# Modeling microbial diversity across different soil ecosystems and environmental covariate selections in tropical SE Asian landscape

**DOI:** 10.1101/2021.04.27.441572

**Authors:** Andri Wibowo

## Abstract

Microbes play essential roles in the ecology of various environments and structure. Diversity of soil microbial communities is an important scientific interest in tropical landscape including in South East Asia. Soil environment in SE Asia is diverse and this influences the microbial diversity. Soil microbes in this study were collected using DNA isolation protocols and the whole microbial community structure of sampled soil was analyzed through next generation sequencing. Soil microbial identification was conducted by amplifying the 16S rRNA gene. Soils were sampled from 4 different environments in Sumatra and Kalimantan ecosystems representing plantation, swamp, peatland, and coastal ecosystems. Measured soil covariates including soil organic carbon C analyzed based on the Walkley-Black method and Kjeldahl method for N. Model of microbes in soil ecosystems were based on Akaike model selection (AIC) by testing the influence of soil covariates on soil microbial phylum. The microbial community was found to be comprising of a total number of 11 phyla. In soil of coastal ecosystem, microbial diversity at phylum level were dominated by Proteobacteria (82.3%), Actinobacteria (9.41%), Acidobacteria (4.7%), and the lowest was Bacteroidetes and Chloroflexi (0.58%). In plantation soil, the microbial abundance order was Firmicutes (41.6%) > Actinobacteria (29.7%) > Acidobacteria (20.0%) > Gemmatimonadetes (5.94%). The soil microbial abundance order in swamp was Proteobacteria (42.2%) > Acidobacteria (20.04%) > Bacteroidetes > (10.46%) > Actinobacteria (9.24%). In soil of peatland ecosystem, taxonomic assignments of microbial at the phylum level were dominated by Firmicutes (66.18%), Proteobacteria(16.94%), and Actinobacteria (16.87%). According to the values of AIC, Firmicutes was a microbial phylum that has high abundance in soil ecosystems influenced by pH covariates with AIC values of −6.54. Other soil covariates show less influence on Firmicutes with AIC values of −5.06. The combination model also show that pH cofactor was the important determinants with AIC values of −0.54 for Firmicutes (pH+C) and Firmicutes (pH+N) models. While AIC value for combination model of Firmicutes (C+N) only equals to 0.14.

## INTRODUCTION

Microbes are the major component of the soil environment. Microbes are ubiquitous in soil environment due to their small size, easy dispersal, capability to exploit a broad range of nutrients and ability to survive well in unfavourable and extreme circumstances. Microorganisms are found in all possible soil environments ranging from those offering ideal conditions for growth and reproduction to extreme environments. Soil environments are one of the environments that harbour population of microbes and are especially suitable for the colonization of microorganisms. These environments vary with respect to the geological features and physicochemical properties including acidic soils. Due to this acidity only microbes can thrive well in this environment which includes mainly bacteria.

Variation of the microbial populations often indicates the change of the soil environment and microbial populations were also influenced by soil environment. Environmental pollution can transform microbial community composition and activity, and there is a dependent relation between microbial diversity and soil environment. Stability of microbial diversity represents the status of a microbial community and this can be used to predict the transformation trend of the environmental quality and soil nutrient conditions. Then microbial community is considered to be one of the most sensitive biological indicators. Soil microorganisms are important component in the cycle of the biological geochemical system and soil microorganism plays an important role in soil self-purification, toxic compound transition, and transformation of the soil environment. Soil microbes are far more sensitive to contaminants than other soil animals and plants and soil microbes can be used as an indicator of the changes in the physical and chemical properties of the soil and environmental quality.

One of versatile method to assess soil microbial community is amplicon next-generation sequencing (NGS) that has been widely used to determine microbial community structures of environmental sampled in fields such as agriculture, ecology, and human health. Microbial species diversity exploration became more attractive due to deep metagenomic sequencing technologies that makes the capacity to sequence multiple samples becomes possible. Diverse ecological interactions between organisms and in biogeochemical processes of nutrient mobilization, decomposition, and gas fluxes determine the microbial communities in soil. Considering this, metagenomic studies of soil communities are very important to understand these processes. In comparison to aquatic environments, physicochemical and biological properties, as well as the presence of compounds that inhibit the polymerase chain reaction makes microbial DNA isolation from soil becomes more challenging. Soil sampling, DNA extraction from microbes in the soil, and data analysis are three important factors need to be considered for a full metagenomic analysis of soils.

South East Asian regions have diverse tropical landscape including the terestrial ecosystems. The soil ecosystems including plantation, swamp, wetland, and coast as these can be observed in Sumatra and Kalimantan ecosystems. This study aims to assess the microbial diversity across different soil ecosystems. This study also measures and models how the soil covariates including pH, C, and N can affect the soil microbial abundances at phylum level. The model was developed based on Akaike (AIC) model selection.

## MATERIALS AND METHODS

### Soil samplings and DNA extraction

Microbes were collected from soils representing 4 different soil environments including plantation, swamp, peatland, and coastal ecosystems of Kalimantan and Sumatra Islands. Each dry soil sample was mixed well and frozen in sterilized tubes at −20 °C until use. Total DNA was independently extracted in triplicate from soil samples using commercially available PowerSoil^®^ DNA. About 0.25 g of soil from each sample was used as starting material. All steps of DNA isolation were conducted following the respective manufacturer’s protocols. A nanodrop spectrophotometer ND 2000 was used to determine DNA concentration and purity of all samples. Following this step, DNA extraction including PCR amplification, purification of PCR products, library preparation, and sequencing, were conducted. Resulted triplicate total DNA samples were barcoded, pooled, and mixed well in one tube.

### PCR amplifications and Next Generation Sequencing

PCR amplifications used Next Generation primer sets attached with Illumina flow cell adapter targeting the 16S rRNA gene of bacteria. PCR amplification was carried out in a total volume of 20 μL containing 40 ng (10 ng/μL) microbial template genomic DNA, 0.6 μL (10 μM) each of forward and reverse primers, and 4.8 μL PCR-grade water. PCR conditions were set at 95 °C for 5 min (initial denaturing step), 30 cycles of 20 s at 98 °C, 20 s at 58 °C, and 30 s at 72 °C, followed by a final extension step at 72 °C for 5 min. Amplicons were quality-tested and size-selected using gel electrophoresis of 1.2% (w/v) agarose concentration and 1× Tris Acetate EDTA run buffer. All PCR was conducted after pooling triplicate samples of total DNA isolates. PCR products were cleaned-up using beads following Illumina MiSeq protocol for amplicon preparation. Library preparation and sequencing were done on an Illumina MiSeq using MiSeq Reagent Kit.

### OTU data analysis

Data analysis was based on the bioinformatics analysis and annotations of the output data were following method by Caporaso et al. (2010) and using QIIME software. QIIME is an open-source bioinformatics tool for performing microbiome analysis from raw sequencing data generated on the Illumina or other sequencing platforms. QIIME gives quality sifting, selecting Operational Taxonomic Units (OTUs), ordered task, phylogenetic remaking, differing qualities investigation and graphical shows. By using QIIME, the information was handled by selecting OTUs based on grouping likeness inside the peruses, and selecting a representative sequence from each OTU. OTU-selecting recognizes exceedingly comparable groupings over the tests and gives a stage for comparisons of community structure. The groupings were clustered into diverse OTUs based on their grouping likeness. Since each OTU was made up of numerous arrangements, a representative sequence arrangement from the OTU was selected for ordered distinguishing proof. The representative sequence for each OTU was classified using classifier and reference databases.

### Soil covariates

Soil covariates that were analyzed in this study were pH, C, and N contents. The determination of soil organic carbon (C) is based on the Walkley-Black (Sahrawat 1982) chromic acid wet oxidation method. Oxidisable matter in the soil was oxidized by 1 N K_2_Cr_2_O_7_ solution. The reaction was assisted by the heat generated when two volumes of H_2_SO_4_ were mixed with one volume of the dichromate. While nitrogen (N) was measured using the Kjeldahl method (Sáez-Plaza et al. 2013), a worldwide standard for calculating the protein content in a wide variety of materials ranging from human and animal food, fertilizer, waste water, and fossil fuels. The method consists essentially of transforming all N in a weighed soil sample into (NH_4_)_2_SO_4_ by digestion with sulfuric acid, alkalizing the solution, and determining the resulting NH_3_ by distilling it into a measured volume of standard acid, the excess of which is determined by titration.

### Statistical analysis and covariate model selections

The obtained microbial abundances from different soil ecosystems were grouping using Principal Component Analysis (PCA). The soil covariates (pH, C, and N contents) then were analyzed using matrix correlation with microbial abundances. While, covariate model selections were developed by testing the soil covariates (pH, C, N) with microbial abundances using Akaike indices (AIC), ΔAIC, and AIC weight. Within this model, pH tests the impact of soil pH on microbial abundance, C tests the impact of soil carbon on microbial abundance, and N tests the impact of soil nitrogen on microbial abundance.

The whole community structure of sampled soil was analyzed through Next Generation Sequencing (NGS) to calculate OTUs. Based on the analysis, the calculated OTUs for all samples were defined by 97% sequence identity among soil microbial representing plantation, swamp, peatland, and coastal ecosystems. NGS analysis indicated that the sampled soil harbours a wealth of different microbial species. The microbial community was found to be comprising of a total number of 11 phyla (Figure 1). In soil of coastal ecosystem, taxonomic assignments of microbial OTUs at the phylum level were dominated by Proteobacteria (82.3%), Actinobacteria (9.41%), Acidobacteria (4.7%), and the lowest was Bacteroidetes and Chloroflexi (0.58%). In plantation soil, the microbial abundance order was Firmicutes (41.6%) > Actinobacteria (29.7%) > Acidobacteria (20.0%) > Gemmatimonadetes (5.94). The soil microbial abundance order in swamp was Proteobacteria (42.2%) > Acidobacteria (20.04%) > Bacteroidetes > (10.46%) > Actinobacteria(9.24%). In soil of peatland ecosystem, taxonomic assignments of microbial OTUs at the phylum level were dominated by Firmicutes (66.18%), Proteobacteria(16.94%), and Actinobacteria (16.87%).

**Figure 1.**
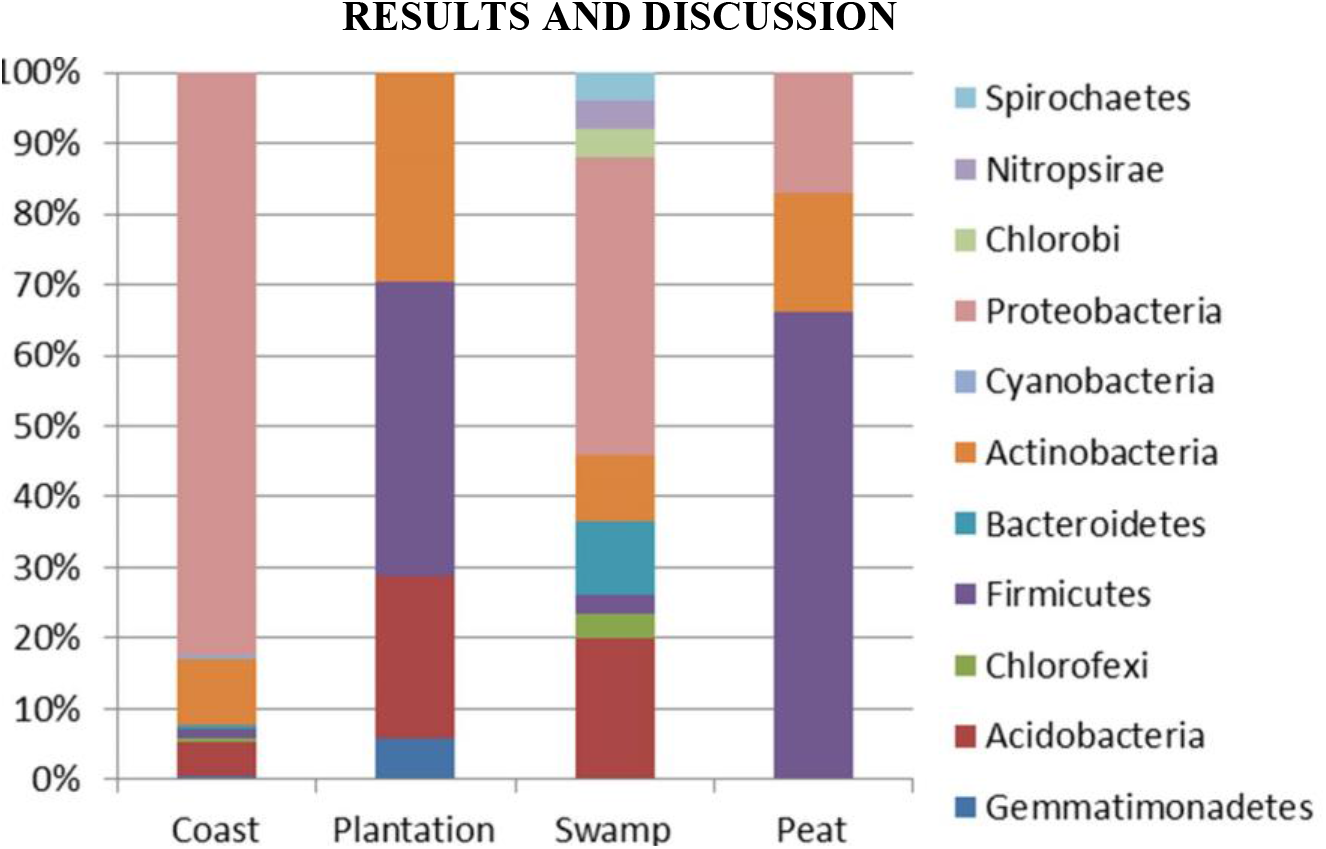
Relative abundance of operational taxonomic units (OTUs) of microbial communities (phyla) from plantation, swamp, peatland, and coastal ecosystems.

Based on the conducted PCA analysis (Figure 2) between microbial communities (phyla) abundance from plantation, swamp, peatland, and coastal ecosystems, there are apparent groupings of microbial phyla. Coastal ecosystem was characterized by abundance of Cyanobacteria with Proteobacteria was performed a distinct clade (Figure 3). Firmicutes, Actinobacteria, Acidobacteria, and Gemmatimonadetes were more related to the peatland and plantation soils. While swamp soils were separated with other soil ecosystems and these ecosystems were characterized by dominations of Proteobacteria that performs distinct clade (Figure 3). OTUs recorded in this study were in agreement with OTUs resulted from other soil microbial studies (Soliman et al. 2017). Soil ecosystems were characterized by maximum representation of members from the phyla like Firmicutes, Proteobacteria, Thermus, Bacteroidetes, and Aquificae (Rawat & Joshi 2019). Kusai & Ayob (2020) have confirmed that the dominant bacterial phyla in peatland were Proteobacteria, Firmicutes, and Actinobacteria. Proteobacteria, Actinobacteria, Acidobacteria, Bacteroidetes and Firmicutes, have frequently been detected as dominant groups in peatland ecosystems.

**Figure 2.**
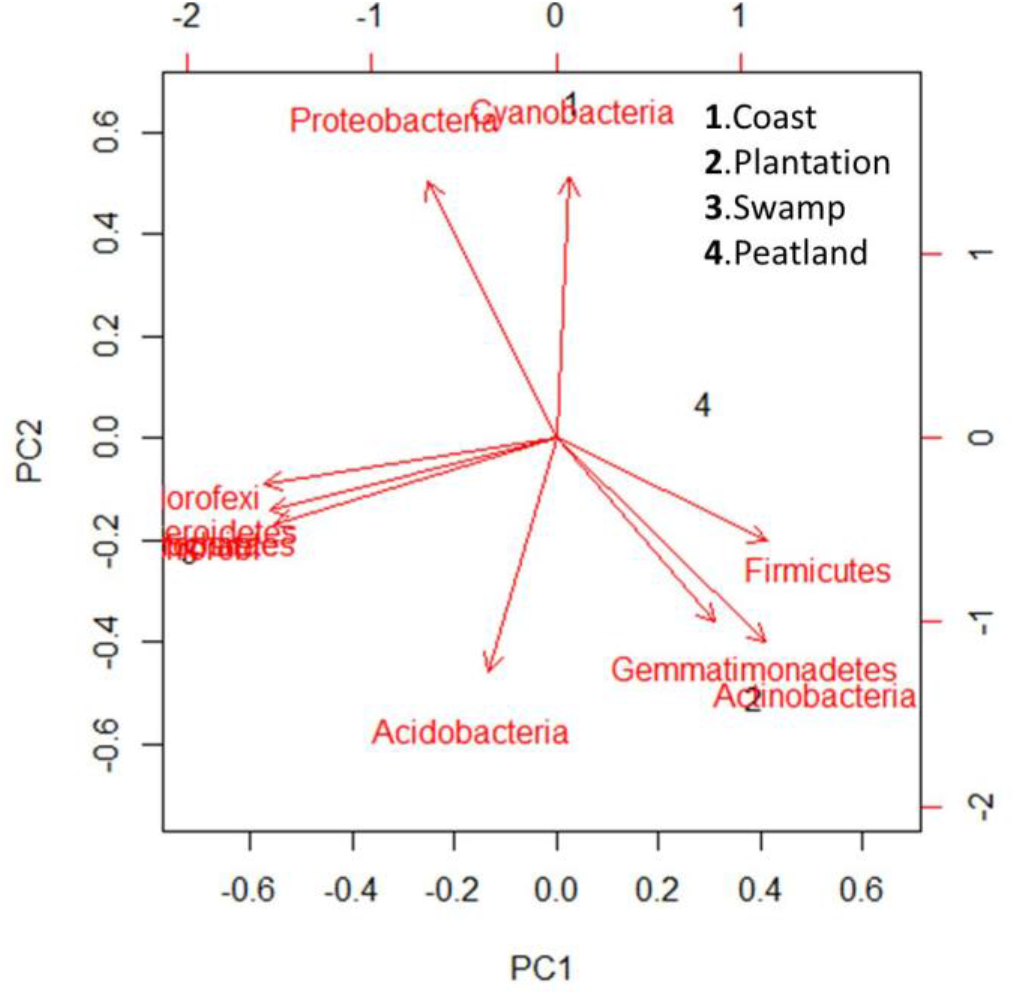
PCA of soil microbial communities (phyla) abundance from plantation, swamp, peatland, and coastal ecosystems.

**Figure 3.**
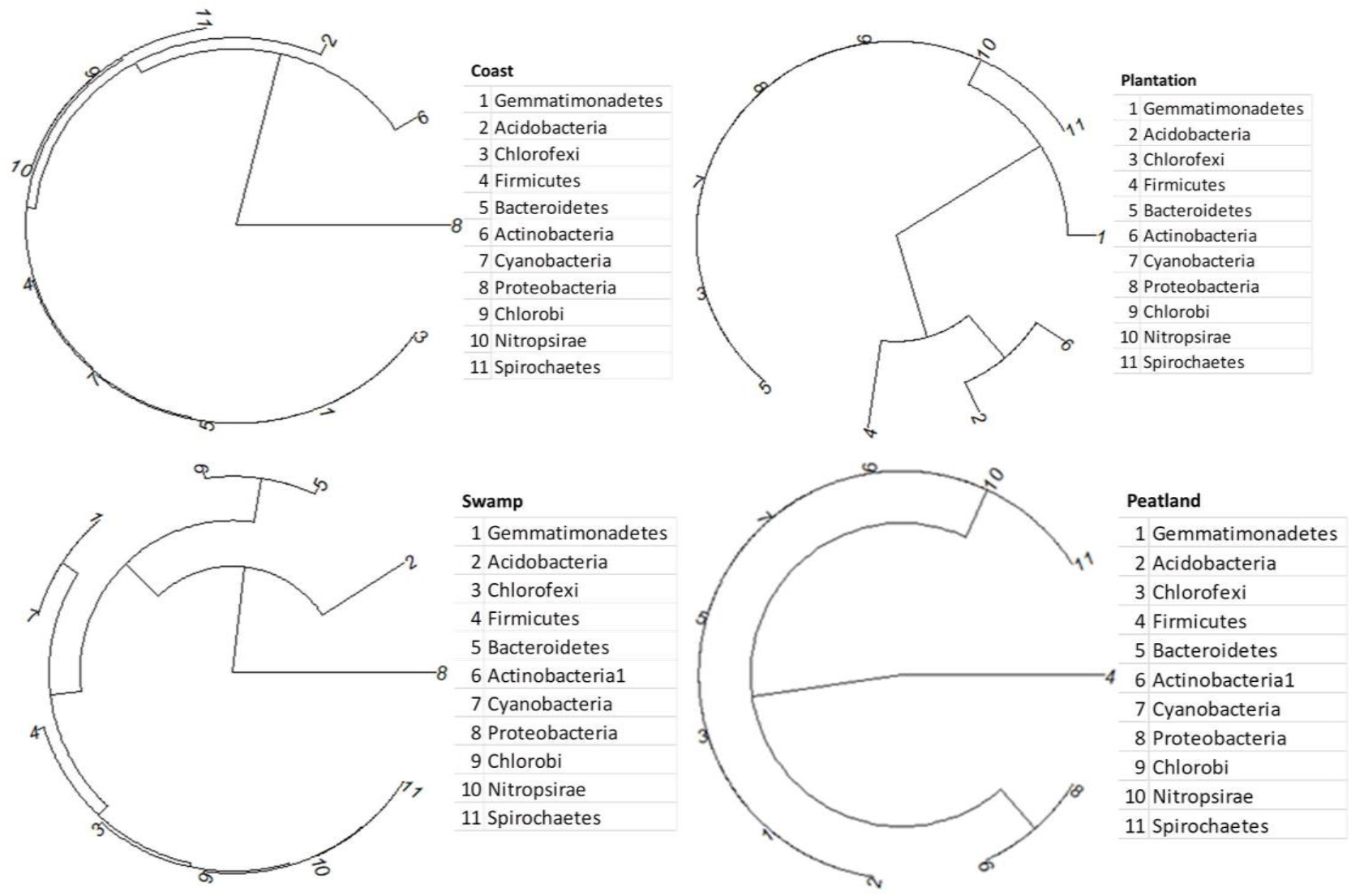
Krona chart of soil microbial communities (phyla) abundance from plantation, swamp, peatland, and coastal ecosystems.

Dominances of some soil microbial phyla were a function of peatland physico-chemical properties, pH, C, and N, and these covariates can cause significant effects on bacterial community structure (Zhou et al. 2017). Figure 4 shows that Firmicutes, a dominant phylum in peatland has significant positive correlation with soil pH and N, and excluding soil carbon. This results agree with numerous previous studies (Qi et al. 2018, Wang et al. 2019). According to the values of AIC (Table 1), Firmicutes was microbial phylum that has high abundance in peatland soil ecosystems influenced by pH covariates with AIC values of −6.54. Other soil covariates show less influence with AIC values of - 5.06. The combination model also show that pH cofactor was the important determinants with AIC values of −0.54 for Firmicutes (pH+C) and Firmicutes (pH+N) models. While AIC for combination model of Firmicutes (C+N) only equals to 0.14. Close relationship of soil pH with bacterial community is due to most of bacteria having rather narrow pH optima and pH might also play an important role in shaping bacterial community. Soil pH has wide association with soil nutrient availability, catabolic activities, soil structure, biomass activities, and mediation of nutrient availability in the soil (Zhalnina et al. 2015). Beside pH, N was also covariate influencing microbial phylum in particular Firmicutes. Li et al. (2016) confirmed that N was significantly positively correlated with Firmicutes (R^2^ = 0.858, P< 0.001). N addition can influence the soil bacterial communities. Zhang et al. (2013) confirmed only N addition significantly (P<0.05) affected soil microbial abundance, richness, and composition.

**Figure 4.**
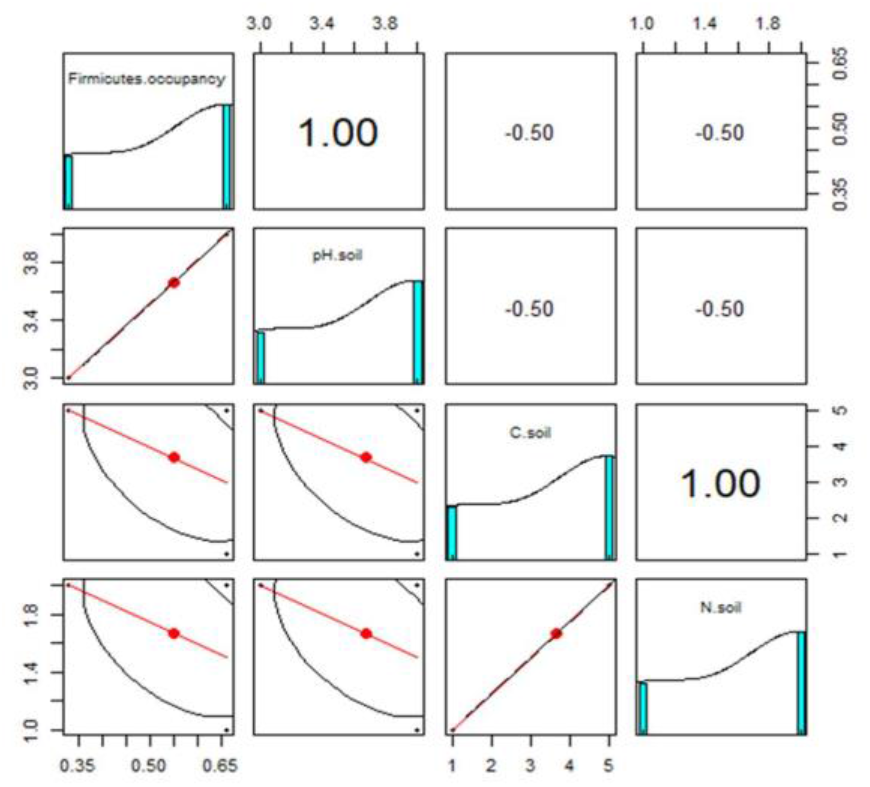
Correlation matrix of Firmicutes phylum abundance with soil covariates (pH, C, N) in peatland soil.

**Table 1.** Firmicutes occupancy models tested for the pH, C, and N covariates (asterisks sign show best model).

| Firmicutes occupancy ($\Psi$) +<br>covariate models | AIC | ΔAIC | AIC<br>weight |
| --- | --- | --- | --- |
| $\Psi(\text{pH})$ | -6.54* | 0.00 | 0.41 |
| $\Psi(\text{C})$ | -5.06 | 0.60 | 0.29 |
| $\Psi(\text{N})$ | -5.06 | 0.65 | 0.29 |
| $\Psi(\text{pH}+\text{C})$ | -0.54* | 0.00 | 0.37 |
| $\Psi(\text{C}+\text{N})$ | 0.14 | 0.68 | 0.26 |
| $\Psi(\text{pH}+\text{N})$ | -0.54 | 0.00 | 0.37 |

In this study, plantation samples were representing disturbed soil environments. In the plantation soil, there were 3 phyla that dominant including Firmicutes, Proteobacteria, and Actinobacteria. This finding is in agreement with other findings and representing the microbal diversity in plantation soil. Liu et al. (2020) confirmed that in disturbed soil the Proteobacteria were the most abundant phylum followed by Acidobacteria. The land use conversion from intact to plantation can disturb soil enviroment and can change its microbial community structure.

